# Sexual isolation maintains species boundaries between Hawaiian crickets in sympatry despite weak habitat isolation

**DOI:** 10.1101/2025.09.04.674279

**Authors:** Raunak Sen, Nicholai M. Hensley, Bhaavya Srivastava, Wout van der Heide, Kerry L. Shaw

## Abstract

Understanding the relative importance of different reproductive isolating barriers in maintaining species boundaries is fundamental to understanding how closely related species coexist without hybridizing. Habitat isolation and sexual isolation are often the first barriers between closely related species, yet they are rarely studied together, making their relative contributions to reproductive isolation unclear. We conducted the first comprehensive study of both barriers in Hawaiian swordtail crickets (*Laupala*), focusing on the recently diverged, sympatric species pair *Laupala kona* and *Laupala hualalai*. Using genomic data, we confirmed that the two species are genetically distinct, which can only be achieved by strong reproductive isolation through a prolonged time period. Through field surveys, we quantified microhabitat use, spatial distribution, and diets, and found extensive ecological overlap. Both species occupied similar substrates, elevations, and microclimates, and occurred in mixed-species aggregations. They only differ in their diets (inferred from stable isotopes), which might have some implications in reducing encounter rates. Despite this weak habitat isolation, they have divergent sexual traits and exhibit strong sexual isolation in lab assays. The species differ significantly in male pulse rate and cuticular hydrocarbon profiles, and females consistently preferred conspecific song stimuli. Mating trials showed almost complete sexual incompatibility between heterospecific pairs, with very rare interspecies copulations. Together, these results demonstrate that sexual isolation, rather than habitat partitioning, maintains species boundaries between *L. hualalai* and *L. kona* in sympatry. Our findings highlight the sufficiency of sexual isolation to prevent hybridization even in the absence of strong ecological divergence, providing rare empirical evidence that sexual barriers alone can maintain reproductive isolation in natural populations.

## 1. Introduction

Species are often defined as reproductively isolated groups, equating the study of speciation to the study of the evolution of reproductive isolating barriers (hereafter RIBs). The evolution of RIBs is important not only for the origin of new species but also to maintain species boundaries when closely related species come into secondary contact. RIBs can be prezygotic or postzygotic depending on the stage in the life cycle of an organism. The first two important prezygotic barriers on secondary contact are habitat isolation and sexual isolation. Often, both habitat and sexual isolation are found between closely related species, which raises some big questions: Is one kind of barrier more important than the other in the evolution and maintenance of reproductive isolation between species? Can they evolve independently of each other? Is either barrier sufficient to prevent interspecies mating? These questions can be asked for any RIBs, but here we focus on the relative contributions of habitat and sexual isolation in preventing interspecies mating, as these are often the first and strongest barriers to gene flow between closely related species (Lackey & Boughman, 2017; Nosil et al., 2024; Shaw et al., 2024).

Habitat isolation is defined as a barrier to gene flow arising from preferences to different habitats. These preferences are based on biological differences between species or inter-species interactions which reduces reproductive encounters between heterospecifics (Coyne & Orr, 2004). We should note that habitat isolation is different from geographic isolation which happens due to the emergence of chance geographical barriers. The adaptation of two races of *Rhagoletis pomonella* to hawthorn and apple trees (Bush, 1969; Jiggins & Bridle, 2004), *Anolis* lizards to different parts of the vegetation (Losos, 2009), and benthic and limnetic stickleback fishes to different depths of the lake (Bell & Foster, 1994), are classic examples of habitat isolation. Sexual isolation is a barrier to gene flow arising from the divergence of prezygotic phenotypes that change reproductive interactions between the sexes (Shaw et al., 2024). Sexual isolation is, on average, the strongest barrier to gene flow in a diverse group of organisms ranging from seed plants (Christie et al., 2022), fungi (Gac & Giraud, 2008), *Drosophila* flies (Yukilevich, 2012) and *Heliconius* butterflies (Garzón-Orduña & Brower, 2018; Mérot et al., 2017; Rosser et al., 2019). But as some authors have noted, these studies have either not quantified the strength of habitat isolation or found it to be equally strong as sexual isolation, raising the question of the relative importance of these two RIBs in reducing gene flow between species (Shaw et al., 2024). Understanding this matters, because we know that the absolute strength of a barrier does not necessarily reflect on its relative strength or importance in reducing gene flow. The relative strength of a barrier depends on the effect of prior acting barriers, in the life cycle of an organism (Dopman et al., 2010; Ramsey et al., 2003). A later acting barrier, even if very strong, would not reduce gene flow to any great degree, if earlier acting barriers had already reduced it substantially. Often sexual isolation acts after habitat isolation, so to understand the relative importance of the two barriers, both should be quantified at the same time.

However, habitat and sexual isolation have been rarely studied together. As mentioned above, most studies of sexual isolation don’t quantify habitat isolation. Similarly, most studies of habitat isolation focus on performance tradeoffs across different habitats and rarely quantify sexual compatibility between these differently adapted species. A recent meta-analysis found that out of 429 papers on habitat isolation published between 2007 and 2022, only 33 papers studied both habitat and sexual isolation (Yukilevich et al., 2024). Among the 33 studies, habitat isolation alone is present in 7, sexual isolation alone is present in 1 and they together act in the remaining 25 cases. However, this dataset has both a poor taxonomic breadth and range of divergence times, which makes drawing any generalizations about the relative importance of both barriers difficult. Support in the theoretical literature is also mixed: while some modelling efforts suggest that sexual isolation alone is insufficient to prevent gene flow without any ecological barriers (Servedio & Bürger, 2014; Weissing et al., 2011), other theoretical treatments find the contrary (Irwin & Schluter, 2022; M’Gonigle et al., 2012). Studying both barriers at the same time is important to not only understand their relative role in keeping closely related contemporary species separate in sympatry but also to throw light on their historical relevance in the speciation process of the group.

Here we investigate the importance of sexual isolation relative to habitat isolation for maintaining species boundaries in the Hawaiian swordtail crickets of the genus *Laupala*. The genus is endemic to Hawaii and contains 38 species inhabiting different islands of the Hawaiian archipelago, but individual species are single island endemic. *Laupala* has the second highest speciation rates in the animal kingdom known to us, only behind African great lake cichlid fishes (Mendelson & Shaw, 2005). Species have historically diverged in allopatry but some closely-related species are found co-occurring in sympatry today (Mendelson & Shaw, 2005; Otte, 1994). Species in this genus are considered morphologically and ecologically cryptic, differing mostly in their sexual signals. They differ in the rhythm of the male courtship song (pulse rate) and in cuticular hydrocarbon (CHC) profiles (long chain carbon compounds present in insect exoskeletons) which are used in chemical communication (Mendelson & Shaw, 2005; Mullen et al., 2007). Male songs and female preferences for the song are species specific, and females strongly prefer conspecific songs over heterospecific songs (Mendelson & Shaw, 2002; Shaw, 2000), suggesting that courtship song acts as a barrier to gene exchange in this radiation. This rapidly speciating group, with minimal ecological divergence but elaborate sexual trait divergence provides a good model clade to understand the role of sexual isolation relative to habitat isolation, in sympatry.

Although the diversity of song and CHC profiles have been documented in many species in the genus, explicit tests of sexual isolation by performing mating trials have not always been done. The most powerful setting to study both habitat and sexual isolation would be where closely related sister species naturally co-occur in sympatry. However, while all *Laupala* species are relatively closely related (likely less than 5 MYA; Mendelson and Shaw 2005), sister species are not known to occur in sympatry. Among some closely related, allopatric species, strong but incomplete sexual isolation has been found (Mendelson & Shaw, 2006; Oh et al., 2013; Shaw & Lugo, 2001). A study of space and habitat use of the more divergent, but sympatric - *L.pruna* and *L.cerasina* showed that the two cricket species are highly mixed both vertically and horizontally, use similar calling sites and exist in mixed-species calling clusters (Xu & Shaw, 2020). Additionally, a documented asynchrony in the diel activities of male singing behavior exists between the two (non-sister) sympatric species *L.paranigra* and *L.cerasina* (Danley et al., 2007). Together these studies suggest that sexual isolation between closely related allopatric species is strong and habitat isolation between divergent sympatric species is weak, but without explicit tests of both on more closely related sympatric taxa, we cannot determine the relative contribution of habitat and sexual isolation to total reproductive isolation.

To fill this gap, we studied sexual isolation and habitat use in the Big Island endemics: *Laupala kona* and *Laupala hualalai*, a naturally co-occurring species pair found in the cloud forests, that diverged less than 0.5 million years ago. First we generated a genome wide SNP dataset to investigate whether they are genetically distinct species with little to no gene flow between them, which would result from strong reproductive isolation. Next, we asked what is the relative contribution of habitat and sexual isolation to total reproductive isolation. Given the importance of sexual trait diversification in the genus and based on the limited previous work on the ecology of species in this genus we hypothesize that reproductive isolation between *L. kona* and *L. hualalai* is mostly driven by sexual isolation, not habitat differences. We conducted surveys in the field and quantified different components of habitat isolation: micro-habitat use, substrate temperature and distance from the ground. We monitored singing activity in the field to estimate temporal overlap/isolation. Shared foraging sites can increase encounter frequency between heterospecifics so we tried to estimate the overlap/ difference in the diets of *L. kona* and *L. hualalai* by using the proxy of stable isotope data. Dietary variation leads to predictable variation in carbon and nitrogen stable isotopes which can be measured from body tissues (Cabana & Rasmussen, 1994; France, 1995; Hobson & Clark, 1992). We collected song data, both in the field and the lab to confirm that they are distinct singing clusters. In *Laupala* and in most singing cricket species, song variation corresponds to distinct breeding populations (Shaw, 1999). Wild collected crickets were brought back to the lab to quantify variation in their CHC profiles. Lab bred progeny of these wild crickets were used in female phonotaxis experiments to test for conspecific song preference. Lastly we did a direct test of sexual isolation by conducting interspecies mating compatibility trials in the lab.

## 2. Methods

### Study site

Our study was based at Makāula ‘O’oma tract (19°43’12” N, 155°56’56” W) of the Honua’ula Forest Reserve on the west coast of the Big Island of Hawaii, where we know *L. kona* and *L. hualalai* exist in sympatry (Otte, 1994; Shaw, 1993). The study site is part of a large tract of cloud forest at elevations above 900 meters on the slopes of Hualālai volcano uphill from the city of Kailua-Kona (see **Figure 1**). The vegetation in the forest is composed of both native and introduced species. Some common native plants were the hapu’u tree fern (*Cibotium glaucum*), the ‘ōhi’a lehua (*Metrosideros polymorpha*) and the ‘ie□ie (*Freycinetia arborea*). Introduced plants included the invasive kāhili ginger (*Hedychium gardnerianum*) and strawberry guava (*Psidium cattleianum*). *Laupala* crickets live mostly in the leaf litter or the forest understory. Where relevant, crickets were collected by netting through vegetation and leaf litter. All field observations were done from February to June 2024.

**Figure 1:**
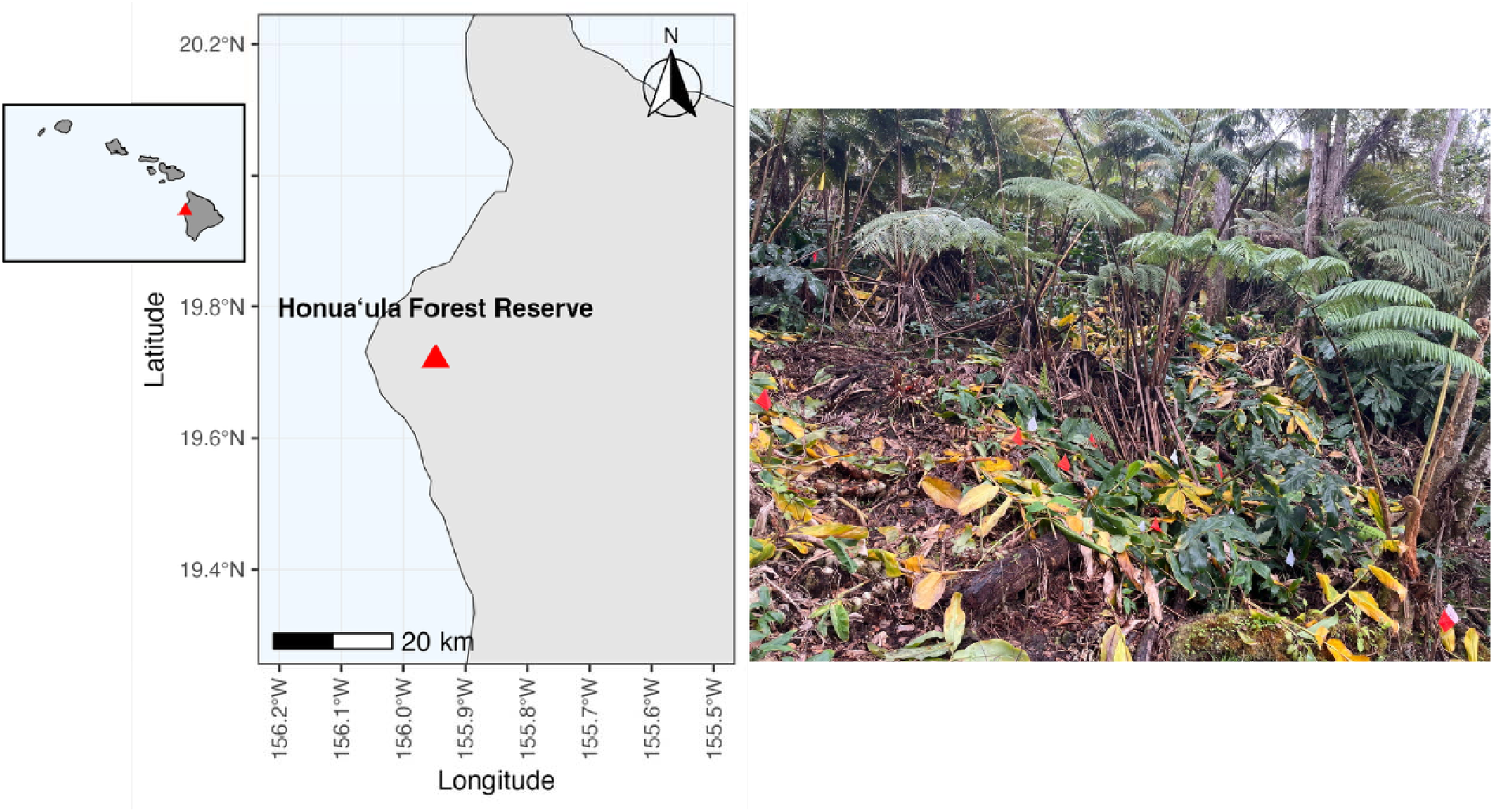
(left) Location of the field site on the Big Island of Hawaii and (right) a representative image of the, habitat of the crickets with the site flags marking where they were heard

#### 2.1 Quantifying population genomic variation between the species

Live crickets were collected from the study site in May 2021, transported back to the lab, phenotyped for song (see Methods section 2.5) and male crickets were assigned species identities. DNA was then extracted from these samples with the QIAGEN DNeasy Blood and Tissue Kit. DNA samples were sent to the University of Wisconsin Biotechnology Center for library preparation and sequencing for Genotype by Sequencing (GBS)(Elshire et al., 2011). Briefly, the enzyme cutters `MspI` and `PstI` were used to digest DNA, which was then ligated to barcodes and common Illumina TruSeq adapters. The ligated DNA was then amplified using PCR and pooled to be sequenced on a shared lane of a Illumina NovaSeq X Plus. 12 samples of *L. kona* and 10 samples of *L. hualalai* were sequenced. Sequence reads were demultiplexed in `Stacks 2.67` (Rochette et al., 2019) and trimmed for adapters and low quality reads with `Trimmomatic v 0.39` (Bolger et al., 2014). Trimmed reads were then aligned to the new version of the *Laupala kohalensis* genome assembly (Hensley et al., in prep) using `bwa-mem2 version 2.2.1` (Li & Durbin, 2009; Vasimuddin et al., 2019). We used `samtools v 1.2` (Li et al., 2009) to convert the files from `sam` to `bam`, sort it and then filter by mapping quality (MQ) score to remove reads with MQ < 20. The filtered `bam` files were used to call variants using `bcftools v 1.20` (Li, 2011) to create a VCF file. We then filtered the VCF file using `vcftools` (Danecek et al., 2011) to remove any loci with more than 20% of the data missing and any in-dels, retaining only single nucleotide polymorphisms (SNPs), retain the SNPs which have a `phred` quality score > 40 (high quality base calls), bi-allelic, have a minimum average read depth of 10 and maximum 40 and a minor allele count > 1. After the filtering steps we retained 65,359 SNPs. We analyzed the SNP dataset in three ways. First, we used a model free approach, namely a principal component analysis (PCA) to summarize the genetic variation between the species. Linkage pruning and PCA calculations were done in `plink v 1.90b7` (Chang et al., 2015; Purcell et al., 2007), that left us with 2236 SNPs. Next, we used a model based approach `ADMIXTURE` (Alexander et al., 2009) to test for the evidence of admixture between *L. kona* and *L. hualalai*. `ADMIXTURE` infers the presence of distinct populations, assigns individuals to populations, identifies admixed individuals and estimates population allele frequencies. Both PCA and ADMIXTURE results were plotted with custom R scripts (adapted and modified from https://speciationgenomics.github.io/). Lastly, we estimated a phylogeny to reconstruct the evolutionary relationship between the two species. A maximum-likelihood phylogeny was estimated in `PhyML 3.0` (Guindon et al., 2010) their “Online execution” platform (http://www.atgc-montpellier.fr/phyml/). The tree was constructed using the HKY85 substitution model with four rate categories and estimated gamma shape parameters. Tree topology was searched using SPR moves starting from a BioNJ tree. A bootstrap analysis with 100 replicates was done to determine node support. 2236 unlinked SNPs were used to build the phylogeny. Since this dataset did not include an outgroup, the tree was rooted at the midpoint of the branch separating the two groups. The topology was annotated and rendered using iTOL v7 (https://itol.embl.de/).

#### 2.2 Quantifying micro-habitat preferences and spatial segregation

After preliminary scouting for areas of sympatry (by listening to calling songs of both species), we chose two 100 m^2^ quadrats at our study site, where we could hear males of both species singing. The crickets in our genus are morphologically cryptic, so we could only locate singing males of either species. Any female crickets, non-adult crickets and non-singing males cannot be assigned species identity in the field, so we only quantify habitat use with singing males. We surveyed the quadrats from 9 to 11 am, when both species sing (see Results section 3.2). We located singing males by ear, listening from different directions and moving nearer to the source of the sound (sensed by higher amplitude). We marked the approximate position of singing males with site flags, using a different colored flag for each species. Two observers were working, together we would each place flags on different halves of a quadrant and then cross check the other half for any singing males missed by the other observer. After placing flags, for each we measured: (1) the substrate temperature using a temperature gun, (2) the elevation from the ground, and (3) categorized the site type as“Leaf litter” or “Above ground vegetation”. We also noted the species of the nearest singing male and measured the distance between them (the nearest neighbor distance; NND) using a transect tape. The NND was classified either as a “conspecific” or a “heterospecific” distance, depending on the identity of the nearest singing male. We collected data on two separate quadrats.

#### 2.3 Quantifying stable isotope variation

Cricket specimens used for C and N stable isotope analysis were frozen within 1-2 hr of their collection in the field. We extracted the CHC’s from individuals and analyzed them to assign species identities based on their distinct CHC profiles. Post CHC extraction, the head and legs were used for stable isotope analysis. We avoided the thorax and abdomen to exclude the digestive tract as recently consumed meals in the gut may bias isotopic values. Samples were dried in an oven at 56°C for 24 hrs before processing. Around 0.4 - 1.2 mg of tissue per sample was weighed out into a 9 x 5 mm tin capsule. Samples were analyzed for C and N content (per cent dry weight) and C and N stable isotope ratios on a Thermo Delta V isotope ratio mass spectrometer (IRMS) interfaced to a NC2500 elemental analyzer at the Cornell Isotope Laboratory (COIL). The primary reference scale used for C was Vienna Pee Dee Belemnite, and the reference for N was atmospheric air. For this analytical sample run the overall standard deviation for the internal animal standard (‘DEER’) was 0.12‰ for ^13^C and 0.10‰ for ^15^N. n=15 *L. hualalai* and n=21 *L. kona* samples were analyzed.

#### 2.4 Quantifying singing activity in the field

We quantified the number of singing males we heard at different time points during the day. We chose two 100 m transects where we could hear both species and marked them every 10 m with a site flag. At each flag, we counted the number of males of each species we could hear within a 1 m radius for 2 min. We repeated this for each of the 10 sites for every hour from 7 am to 7 pm, on the same day, totaling 13 time points. The two transects were done on separate days. For analysis, we summed up the total number of singing males we heard of each species (across 10 sites) at each time point. To analyze cricket singing activity, we fitted a linear mixed-effects model using the `lmer` function from the `lme4` (Bates et al., 2015) package in R. The number of singing males served as the response variable, with species (*L. kona* vs. *L. hualalai*) and time of the day (hour from 7am-7pm) as fixed effects, including their interaction to test for species-specific temporal patterns. Transect was included as a random intercept term to account for baseline differences between sampling locations and the non-independence of repeated hourly measurements within each transect. Model assumptions were assessed through visual inspection of residual plots. Effect-sizes of the model were calculated using the `eta_squared` function from the `effectsize` package in R.

#### 2.5 Quantifying pulse rate variation

To document song divergence between the two species, we quantified the song pulse rate variation in males of either species, both in the wild and in the lab. As lab conditions are more stable than the field (i.e. without fluctuating temperatures), we quantified song differences between the two species in both the lab and field to assess if any observed differences in the wild were consistent in a controlled lab environment. In May 2023, we recorded singing males in the wild using a Sony digital voice recorder (Model: ICDUX570BLK) with the microphone attached to the end of a dowel. We probed the vegetation and leaf litter until we clearly heard a singing male or a pair of singing males and then recorded their songs. Lab recorded crickets were initially collected in the wild as nymphs in May 2021. They were reared in the lab in 4 quart plastic canisters lined with moist Kimwipe tissues on a diet of Organix organic cat chow. On reaching maturity they were separated by sex. Males were placed in individual cups, and their songs were recorded using Olympus digital voice recorders (Model: VN 722PC) in a temperature-controlled room (19-21 C). *Laupala* crickets have a simple song structure which is a train of pulses. We analyzed the song files in Raven Pro 1.6.5 (https://www.ravensoundsoftware.com/) by calculating the pulse period, which is the time between the beginning of one pulse and the next. We took 5 independent measurements of pulse period and averaged them for the lab recorded songs and 3 independent measurements for the wild recorded ones. We calculated the pulse rate by using the inverse of the pulse period. A histogram of the distribution of pulse rates was plotted for songs recorded in the wild (n=17 *L. hualalai* and n=20 *L. kona*) and in the lab (n=17 *L. hualalai* and n=28 *L. kona*).

#### 2.6 Quantifying CHC variation

CHCs were extracted from previously frozen crickets, which were collected in the wild in May 2023. To extract the chemicals, a cricket was thawed and washed for 5 minutes in 300 μl of hexane in a glass vial that also contained 0.034 g each of nonadecane (C19 alkane) and pentatriacontane (C35 alkane) as internal standards for the chromatograph (Stamps & Shaw, 2019). The resulting extract was then filtered through glass wool into a clean glass vial. After filtration, the extract was evaporated under a gentle nitrogen stream and stored frozen at −20 °C.

When all CHC samples were extracted and ready to be run on the chromatograph, the extracts were resuspended in 60 μl of clean hexane with no internal standards and the solution was pipetted into a 100 μl glass insert fitted inside the autosampler vial. A Shimadzu AOC-20i autoinjector was used to inject 1μl of the samples into a Shimadzu GC-2014 gas chromatograph with flame ionization detection (GC-FID) equipped with an HP-5 capillary column (length: 20 m, diameter: 0.180 mm, film thickness: 0.18 μm). The injection port temperature was 300°C. The inlet pressure was 144.2 kPa and column flow rate was 0.8 ml/min for a linear velocity of 32.5 cm/s. The oven temperature was held at 60°C for one minute and ramped to 200°C at a rate of 20°C per minute. The rate was then decreased to 5°C per minute until the oven temperature reached 320°C, where it was held for 15 minutes. The total program time per sample was 47 minutes. Peaks were integrated in Shimadzu’s LabSolutions software, using the autointegrate function with a slope of 18,000 μV/s. Integrated peak tables were exported from LabSolutions to Microsoft Excel and R for further analysis.

Peak tables were imported into RStudio as data frames. Percent abundance for each peak was transformed using a centered-log ratio transformation (Aitchison, 1982; Mullen et al., 2007).

Peaks were filtered so that only those with a percent abundance ≥1 remained. The prepared peak tables were then aligned using the package `GCalignR` (Ottensmann et al., 2018). The resulting alignments were used in a principal components analysis (PCA) using a covariance matrix, using the function `prcomp`. To visualize the variation in CHC space, PC1 and PC2 were plotted.

#### 2.7 Quantifying female acoustic preferences

Phonotaxis trials were used to quantify female acoustic preferences. Laboratory bred and raised females of both species were used in the experiment after they reached sexual maturity (> 14 days post final molt). Playback experiments with two song stimuli were run in a 47 cm circular arena placed within a temperature controlled (20+1°C) anechoic chamber (Acoustic systems). Songs were digitally generated and played from two 8.5 cm speakers (Radio Shack model 40–1218) placed 180◦ apart just outside the arena. From each speaker, a pulsed, sinusoidal tone was played back through a 16-bit digital/analogue converter (Tucker-Davis Technologies, Gainesville,FL). The pulse amplitude envelope had a rise time of 10 ms and a fall time of 30 ms and the acoustic output was filtered at 10 kHz using a Krohn-Hite filter (model 3322) to avoid aliasing. Sound pressure levels were equalized to 90 dB on a 4.0-pulses per second (pps) tone monitored with a Bruel and Kjaer sound pressure level meter (type 2230, fast-root-mean-square setting). Songs with two pulse rates were used: 1.4 pps (mean pulse rate of *L. hualalai* in the lab) and 2.1 pps (mean pulse rate of *L. kona* in the lab), but pulse duration and carrier frequency were kept constant at 40 ms and 5.0 kHz respectively.

For the trial, a female was placed in a plastic cup in the center of the arena. The two songs were broadcast simultaneously from either speaker for 5 min. Which speaker played which song was randomized. The cup was remotely raised following the 5 min, giving females access to the arena. A ‘response’ was recorded if the female entered a predetermined area 10 cm from either speaker (for a schematic of the arena, see Figure 7 and Shaw & Herlihy, 2000). A ‘no response’ was recorded if the female did not enter either area in 3 minutes. All trials were conducted between 10 am and 4 pm. A total of n=22 *L. hualalai* and n=25 *L. kona* females were trialled in the phonotaxis chamber.

#### 2.8 Quantifying inter-species mating (in)compatibility

No-choice mating trials were done to quantify mating (in)compatibility. The crickets used in the experiments are lab-bred progeny (F2 and F3) of wild caught females. Young nymphs (∼ 3 months post hatching) were separated by sex. Around 5 months post-hatching, older nymphs were separated into individual specimen cups lined with a moist kimwipe and monitored every other day for maturation (final molt). All crickets were kept in a temperature-controlled room set to 20+1°C and a 12 hr light dark cycle. All individuals were used between 21 to 30 days post maturity as this age is known to be the window of prime sexual receptivity. This experiment was partly done in Feb - March 2024 and in March - May 2025. There was no difference in the results at two time points, so we combined the two datasets.

The mating experiments consisted of three stages: a pre-trial, the test trial followed by a post-trial. The pre-trial is a sexual receptive test to ensure that the crickets that go into the experiment are sexually mature. Conspecific male and female pairs were put in a petri dish and their mating interactions were observed. *Laupala* crickets have a long and protracted courtship ritual lasting 5-7 hrs involving the male producing and transferring many spermless micro spermatophores (hereafter called “micros”) followed by the sperm filled macro spermatophore (hereafter “macro”) (Shaw & Khine, 2004). Pairs passed the sexual receptivity test if the male produced a micro and both male and female were in a transfer position, poised to transfer the micro. The trial was ended at this point (before any micro transfer). The test-trial happened after 1-2 days when the sexually receptive crickets were paired with heterospecifics of the opposite sex. 1-2 days after the test trial, they were paired with conspecifics for the post-trial, but this time they were left alone to mate. For all test and post trials, a male and a female cricket were put together in a closed Petri dish with a moist kimwipe and their interactions were video recorded with a Sony HDRCX405 Camcorder. Trials were started around 11:30 am and recorded for ∼7 hrs in a temperature-controlled room set to 20+1°C. Trials ended after 7 hrs or until the fate of a produced macro was known. The videos were then manually scored for different behaviors. For each trial, we scored four behaviors: whether the male produced and/or transferred any micro(s), and if the male produced and/or transferred a macro. A successful micro transfer was defined as courtship success and a successful macro transfer was defined as mating success. 74 individuals were tested in test and post trials. To analyze courtship and mating success, we fitted a generalized linear mixed-effects model using the `glmer` function from the `lme4` (Bates et al., 2015) package in R. Courtship/ mating success was the response variable, with species (*L. kona* or *L. hualalai*), sex (male or female) and pairing type (heterospecific or conspecific) as fixed effects. Individual identity was included as a random intercept term to account for individual differences in sexual receptivity. Model significance was assessed using Type III Wald chi-square tests. Model assumptions were assessed through visual inspection of residual plots. Predicted probabilities of courtship and odds ratio for each pairing type were estimated from the model using estimated marginal means with the `emmeans` package (Lenth et al., 2025). The data was also grouped by pairing groups (four combinations including males and females of each species) and plotted in **Figure 9** for easier visualization.

## 3. Results

All statistical analyses were done in RStudio version 2024.09.1+394.

### 3.1 Population genomic variation

In all the population structure analyses, individual samples are consistently associated with their respective species. The PCA neatly groups the samples into two separate species clusters with PC1 explaining 55.8% of the total genomic variation (**Figure 2a**). In the ADMIXTURE analysis, the genetic data was best explained by two distinct genetic clusters (K=2), with a few *L. kona* individuals showing low levels of admixture (**Figure 2b**). The maximum-likelihood phylogeny supports the distinct monophyly of the two species (**Figure 2c**), in line with the other two analyses.

**Figure 2:**
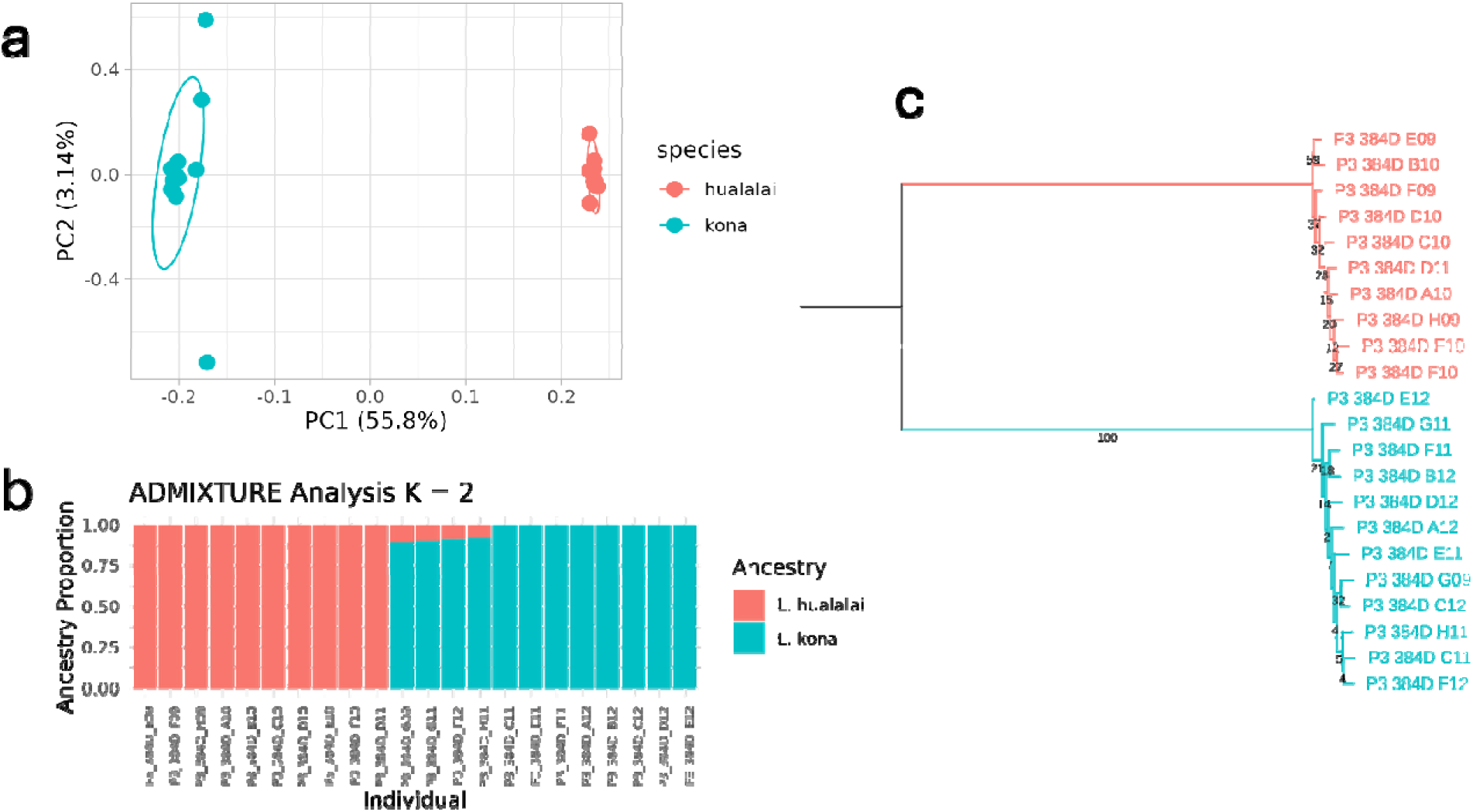
Population structure of n=12 *L. kona* and n= 10 *L. hualalai* individuals (a) Principal components analyses of 2236 genome-wide SNPs. PC axes 1 and 2 explain 55.8% and 3.14% of the variation, respectively. (b) ADMIXTURE analysis with the most likely number of clusters (K=2). Each vertical bar represents an individual which is partitioned into two colors representing distinct ancestries. (c) A phylogenetic tree of all individuals which was rooted at the midpoint of the branch separating the two monophyletic species groups

### 3.2 Micro-habitat preferences and spatial segregation

The number of singing males of either species we recorded is given in **Table 1**.

**Table 1:** Number of singing males of either species in the two quadrats.

| Quadrat | Area(m<sup>2</sup>) | No. of *L. hualalai* males | No. of *L. kona* males |
| --- | --- | --- | --- |
| 1 | 100 | 15 | 6 |
| 2 | 100 | 18 | 19 |

The plots were analyzed jointly as preliminary analysis of them separately yielded similar results. Crickets of both species occupied micro-habitats with similar substrate temperatures (**Figure 3a**, Welch two sample t-test, t = −0.16225, df = 43.998, p-value = 0.8719) and there was no significant difference between the two species in vertical stratification (measured by elevation from the ground, **Figure 3b**, Wilcoxon rank sum test, W = 483, p-value = 0.1717).

**Figure 3:**
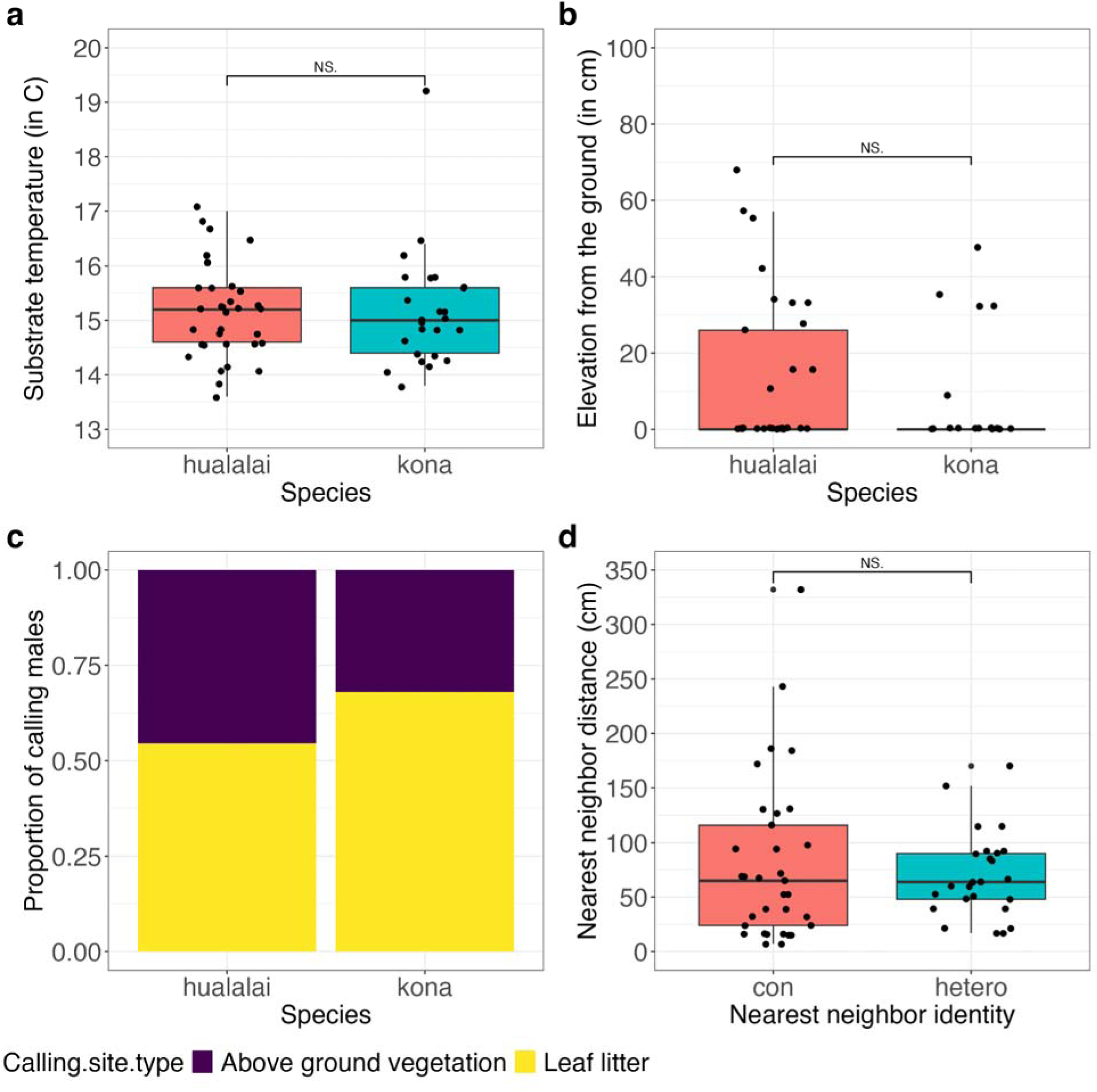
Comparison of (a) substrate temperature of micro-habitats and (b) elevation from the ground of singing males of both species *L. hualalai* and *L. kona* (c) Calling site preferences of both species and (d) Nearest neighbor distances compared between conspecific and heterospecific neighbors

*L. hualalai* and *L. kona* also did not differ in their use of calling sites (□^2^ = 0.587, p-value = 0.4435), with a slightly higher proportion of crickets of either species singing from the ground leaf litter compared to above ground vegetation (**Figure 3c**). Singing clusters consisted of both species: nearest neighbor distance (NND) did not differ significantly between conspecific or heterospecific neighbors (**Figure 3d**, Wilcoxon rank sum test, W = 398, p-value = 0.8259).

### 3.3 Stable isotope variation

There is a huge degree of overlap between the 95% confidence ellipses for the isotopic values of *L. hualalai* and *L. kona* in the two-dimensional δ^13^C and δ^15^N space (**Figure 4**). *L. hualalai* occupies a subset of the isotopic space covered by *L. kona*, although their sample size is a bit lower (n=15) than that of L. kona (n=21). There is no significant difference between the δ^15^N values of both species (t = −1.4641, df = 33.983, p-value = 0.1524). However, *L. kona* had significantly higher δ^13^C values than *L. hualalai* (t = −2.7623, df = 33.918, p-value = 0.0092).

**Figure 4:**
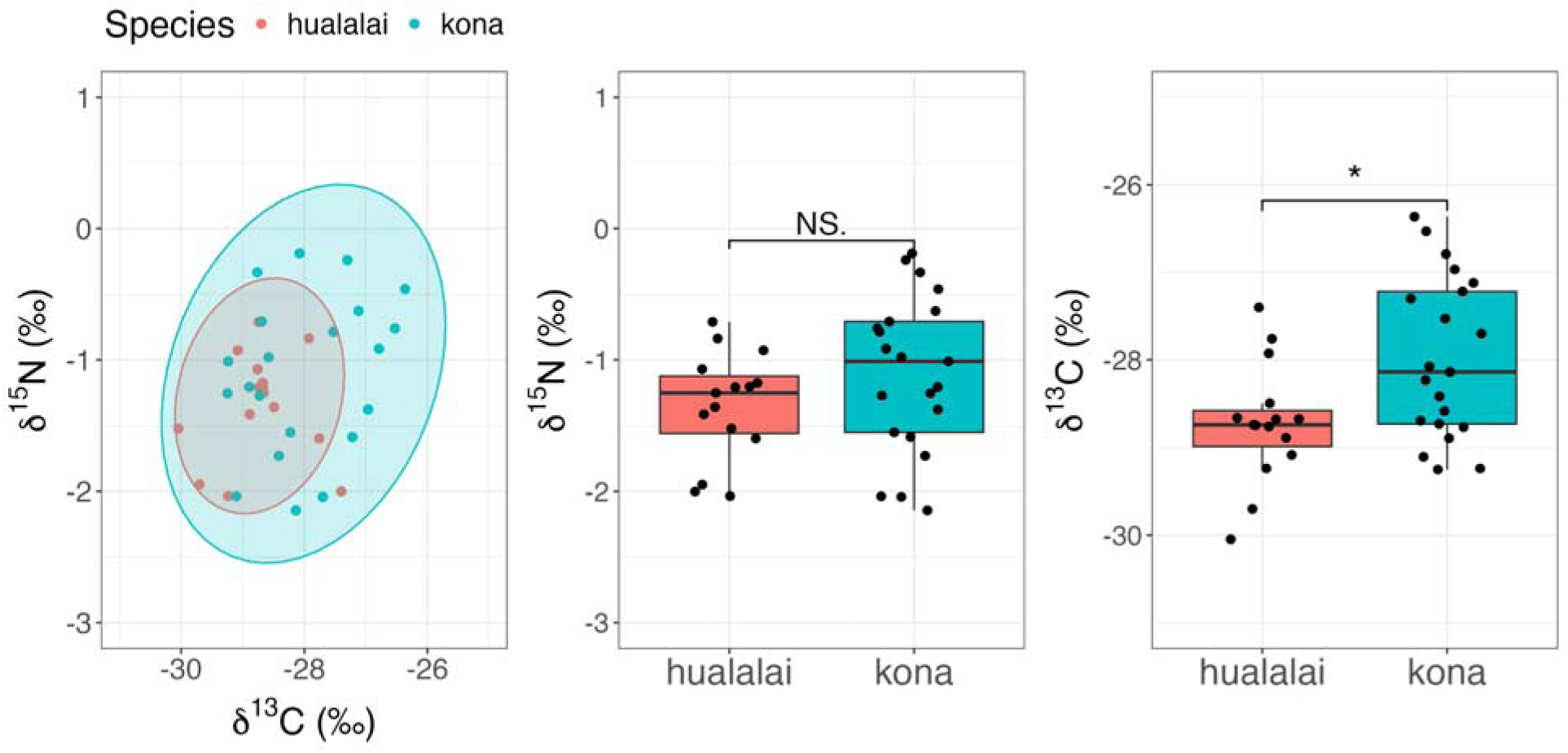
δ^13^C and δ^15^N variation between *L. hualalai* and *L. kona*

### 3.4 Singing activity in the field

*L. hualalai* sang predominantly earlier in the day, with peak singing activity around 9-10 am and a minor spike later in the evening around 7 pm. Whereas peak singing activity of *L. kona* was in the afternoon from 1-3 pm (**Figure 4**). At both transects, we had more *L. kona* males singing than *L. hualalai* males. Linear mixed-effects modeling revealed significant differences in cricket singing activity between *L. kona* and *L. hualalai* (F□,□□ = 21.23, p < 0.001). Both species exhibited temporal variation throughout the day (F□□,□□ = 3.77, p = 0.002), but with significantly different daily patterns (interaction: F□□,□□ = 5.45, p < 0.001), with *L. hualalai* singing earlier in the day compared to *L. kona*. `Species` explained 45% of the variance in cricket singing activity (η²p = 0.45), while the `Species × Time of day` interaction accounted for 72% of the variance (η²p = 0.72), indicating strong species-specific temporal patterns.

### 3.3 Pulse rate variation in the wild and in the lab

Analysis of pulse rate distribution of songs collected in the wild revealed a bimodal distribution of pulse rates, the two clusters corresponding to the two species *L. hualalai* and *L. kona* (**Figure 5**, top panel). We get a similar result from lab recorded songs of crickets collected in the wild (**Figure 6**, bottom panel). *L. kona* sings at a significantly faster pulse rate (see **Table 2**) than *L. hualalai* either in the wild (*L. kona* 1.89 ± 0.066; *L. hualalai* 1.11 ± 0.048) or the lab (*L. kona* 2.17 ± 0.096; *L. hualalai* 1.36 ± 0.043). Pulse rates are significantly faster in the lab compared to the wild, for both species (see **Table 2**).

**Figure 5:**
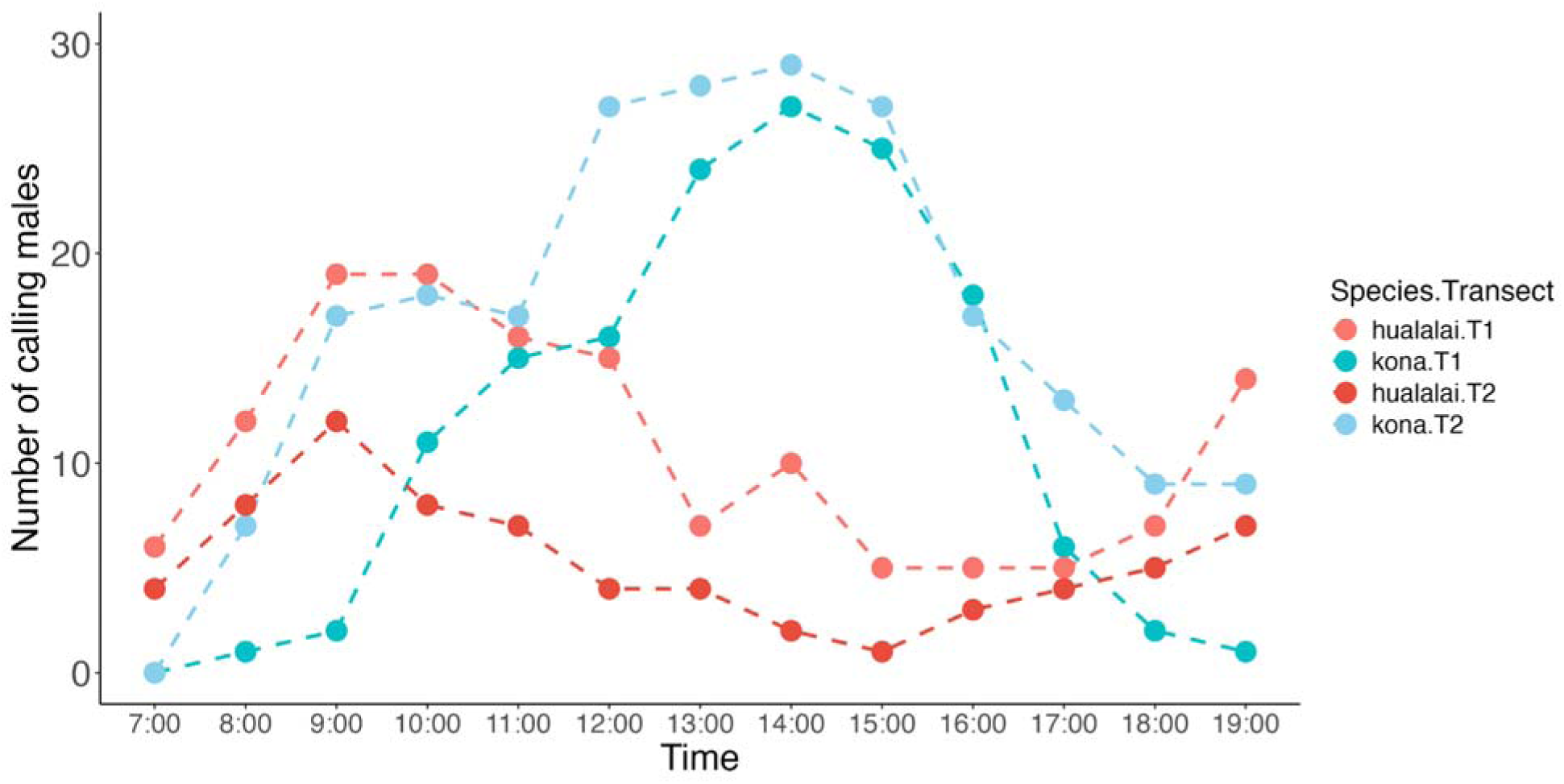
Diel pattern of male singing behavior of *L. hualalai* and *L. kona* across two transects. The dots are the total number of singing males we heard of each species (summed across 10 sites) at each time point, connected by imaginary lines.

**Figure 6:**
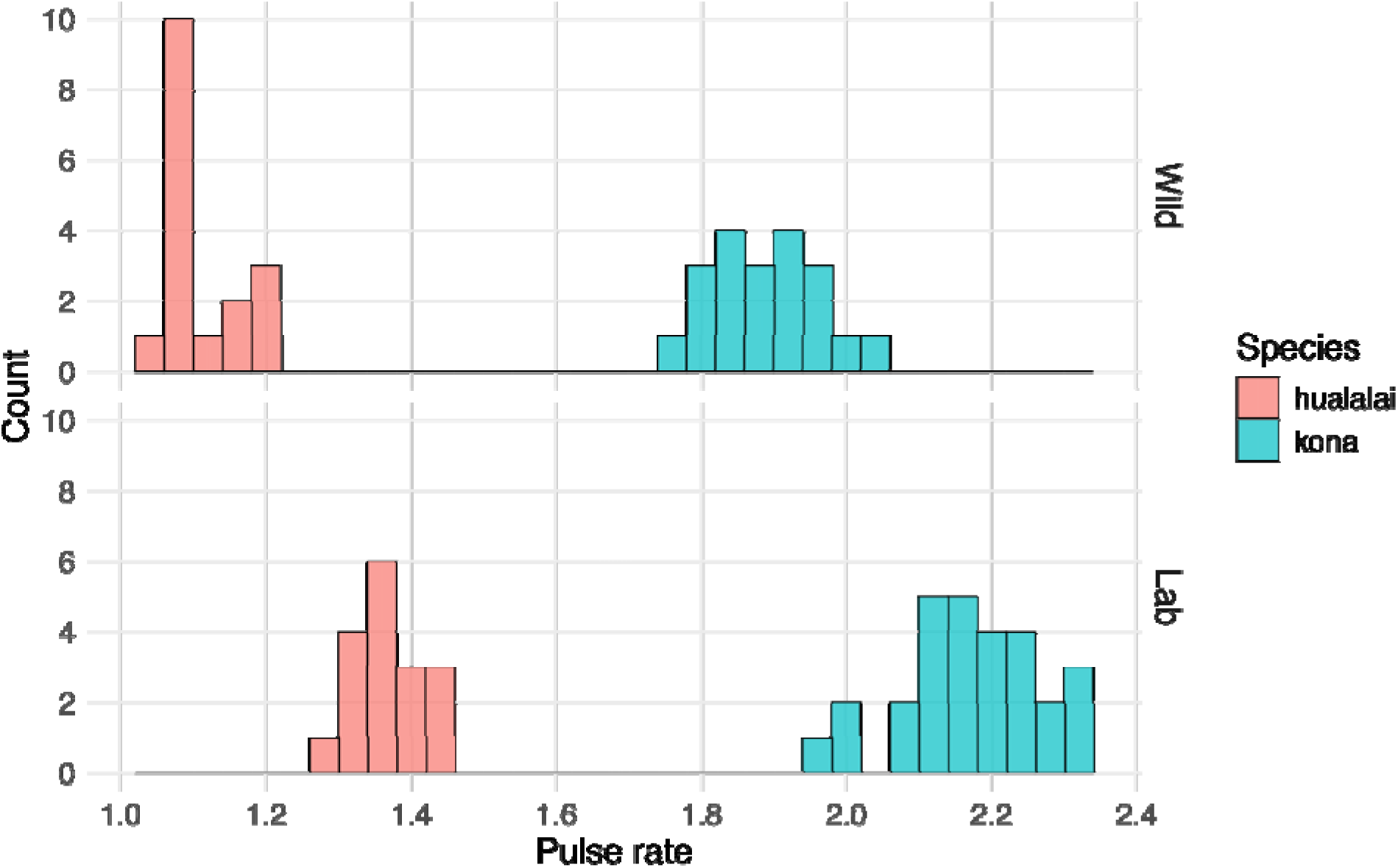
Pulse rate distribution of *L. hualalai* and *L. kona* analyzed from songs collected in the wild (top) and the lab (bottom)

**Table 2:** Welch two sample t-test results for all combinations.

| Comparison | t-statistic | p-value | Signifiant ? |
| --- | --- | --- | --- |
| *L. kona* vs *L. hualalai* (wild) | -41.47 | 7.68e-31 | yes |
| *L. kona* vs *L. hualalai* (lab) | -38.74 | 1.25e-33 | yes |
| *L. hualalai* (wild vs lab) | -16.288 | 5.96e-17 | yes |
| *L. kona* (wild vs lab) | -12.235 | 4.62e-16 | yes |

### 3.4 CHC variation

After pulse rates (song), CHCs are the next important axis of sexual trait diversification in *Laupala*. We looked at the differences in CHC profiles of *L. hualalai* and *L. kona*. A representative chromatogram of either species is shown in **Figure 7a**. Qualitatively, from the chromatograms we can tell that *L. hualalai* has a greater number of peaks than *L. kona*. *L. hualalai* also has some peaks with later elution times, signifying relatively longer chain CHCs, which are missing in *L. kona.*, Similarly, *L. kona* has some peaks with lower elution times, indicating relatively shorter chain CHCs, which are missing in *L. hualalai*. 42 peaks were retained after aligning all the peaks across all samples (n = 24 *L. hualalai* and n = 27 *L. kona* samples). A principal components analysis (PCA) on the dataset resulted in 5 principal components (PCs) with eigenvalues greater than 1, which explained 91.27% of the total CHC variation. A MANOVA found significant difference between the species (Wilks’ λ = 0.05, *F* = 169.51(5,45), *p* < 2.2*10^-16^), driven primarily by PC1 and PC2 (univariate ANOVA, PC1: *F* = 44.18, *p* = 2.34*10^-8^; PC2: *F* = 644.71, *p* < 2.2*10^-16^). PC1 and PC2 together explain 83.2% of the total CHC variation and neatly cluster the data into two species groups (**Figure 7b**). Using PC1 and PC2, a linear discriminant analysis (LDA) with jackknifed cross-validation correctly classified 23/24 of the *L. hualalai* samples and all the *L. kona* samples, representing a predictive accuracy of 98.04%.

**Figure 7:**
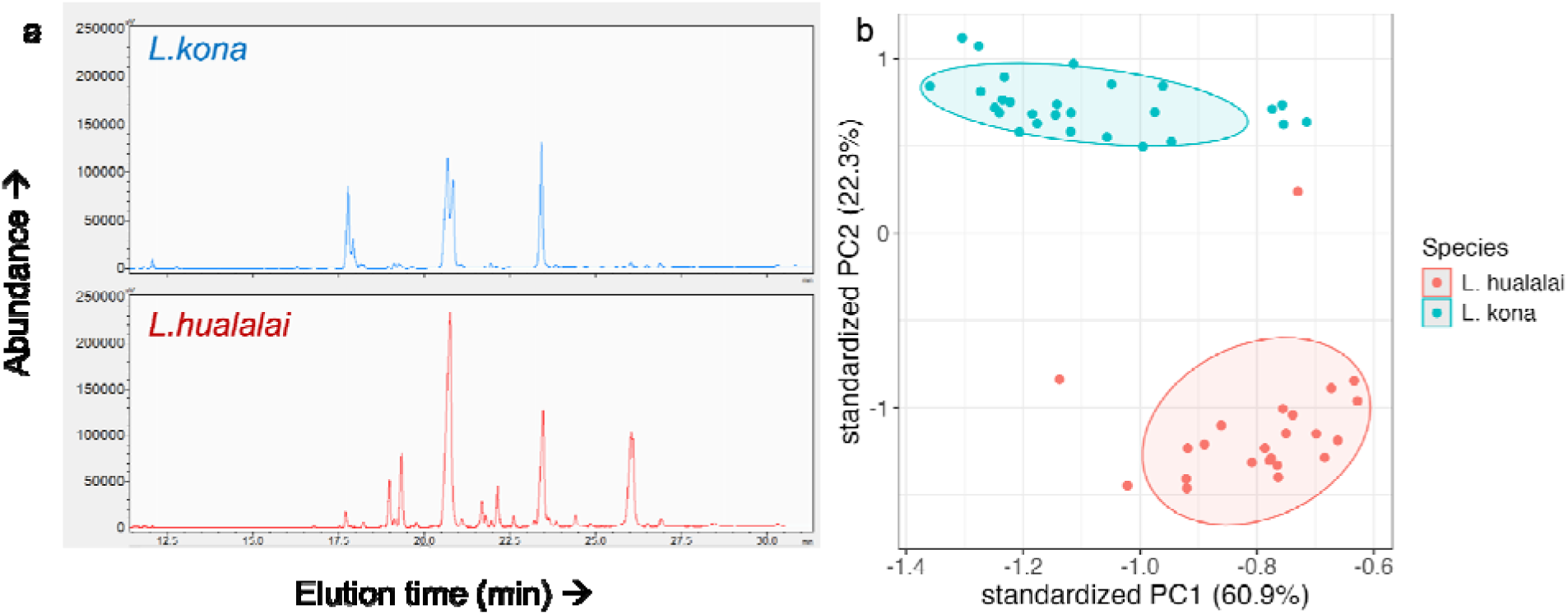
(a) Representative chromatograms of *L. kona* (top) and *L. hualalai* (bottom) (b) PCA plot of the CHC variation in *L. hualalai* and *L. kona*; PC1 and PC2 explain 60.9% and 22.3% of the CHC variation

### 3.5 Female acoustic preferences

*L. kona* females were more responsive to the phonotaxis trials than *L. hualalai* females (Chi-squared test, □^2^ (2) = 29.99, p-value = 3.075e-07). Among the females which were responsive, *L. hualalai* and *L. kona* always chose the slow (1.4 pps) and fast (2.1 pps) pulse rates respectively, which corresponds to their conspecific pulse rates in the lab (Chi-squared test, □^2^ (2) = 29.99, p-value = 3.075e-07) (**Figure 8**).

**Figure 8:**
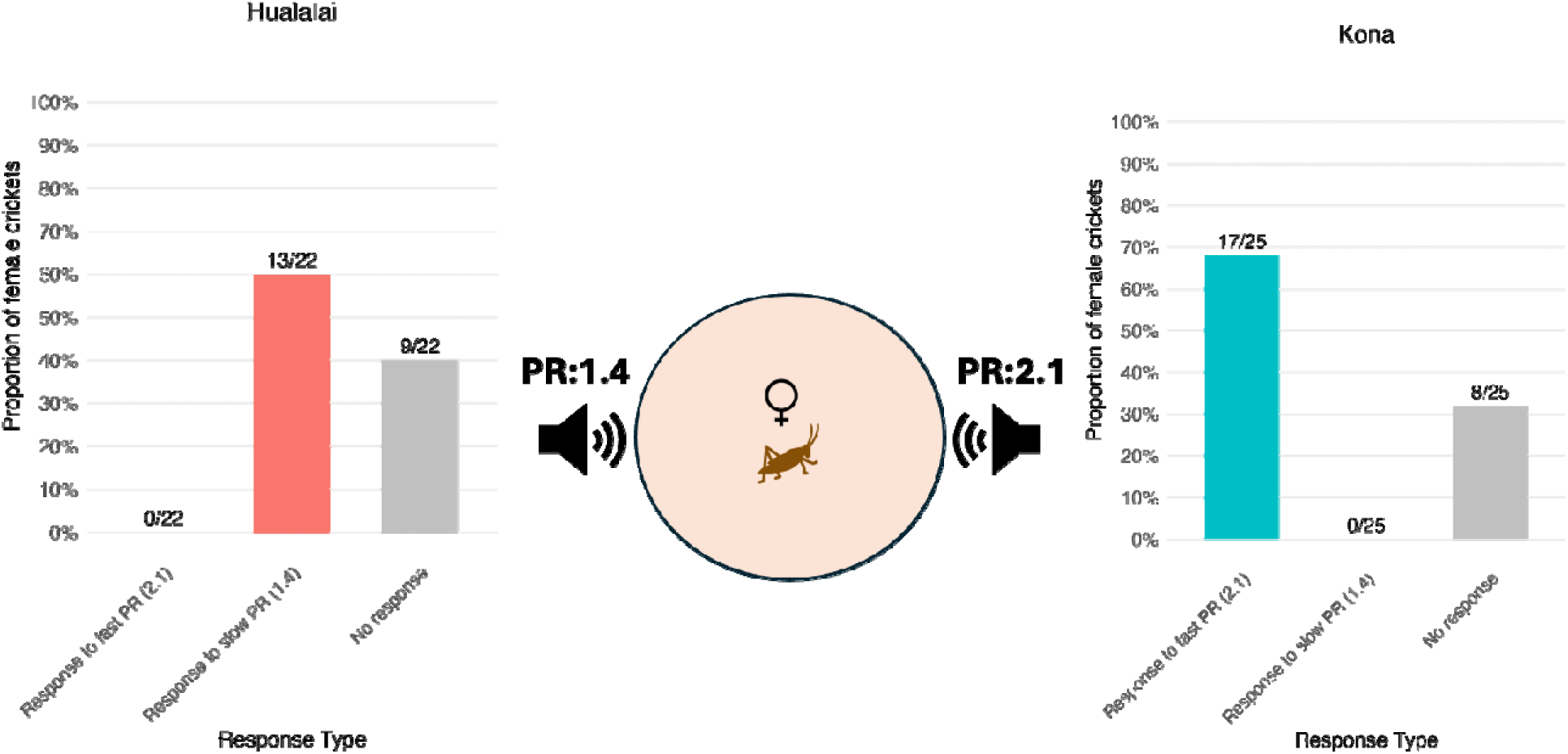
Proportion of female crickets of *L. hualalai* (left) and *L. kona* (right) that responded to either the fast (2.1 pps) or slow (1.4 pps) pulse rate or did not respond to either, with a schematic of the phonotaxis arena in the center. The pulse rate values were chosen based on the mean pulse rate of either species in the lab.

### 3.6 Inter-species mating trials

Courtship and mating was higher in the conspecific pairs, compared to the heterospecific pairs (**Figure 9**). `Pairing type` (heterospecific vs conspecific) was a significant predictor of both courtship (χ² = 45.30, df = 1, p < 0.001) and mating success (χ² = 16.29, df = 1, p < 0.001). Neither `Species` nor `Sex` significantly affected courtship behavior (`Species:` χ² = 1.78, df = 1, p = 0.18; `Sex:` χ² < 0.01, df = 1, p = 0.99) or mating outcomes (`Species:` χ² = 1.24, df = 1, p = 0.26; `Sex:` χ² = 0.01, df = 1, p = 0.91). The odds of courtship and mating were 55 and 73.2 times higher in conspecific pairings. Inter-species matings were extremely rare: 1/37 of the heterospecific pairs mated, signifying strong sexual isolation between the two species.

**Figure 9:**
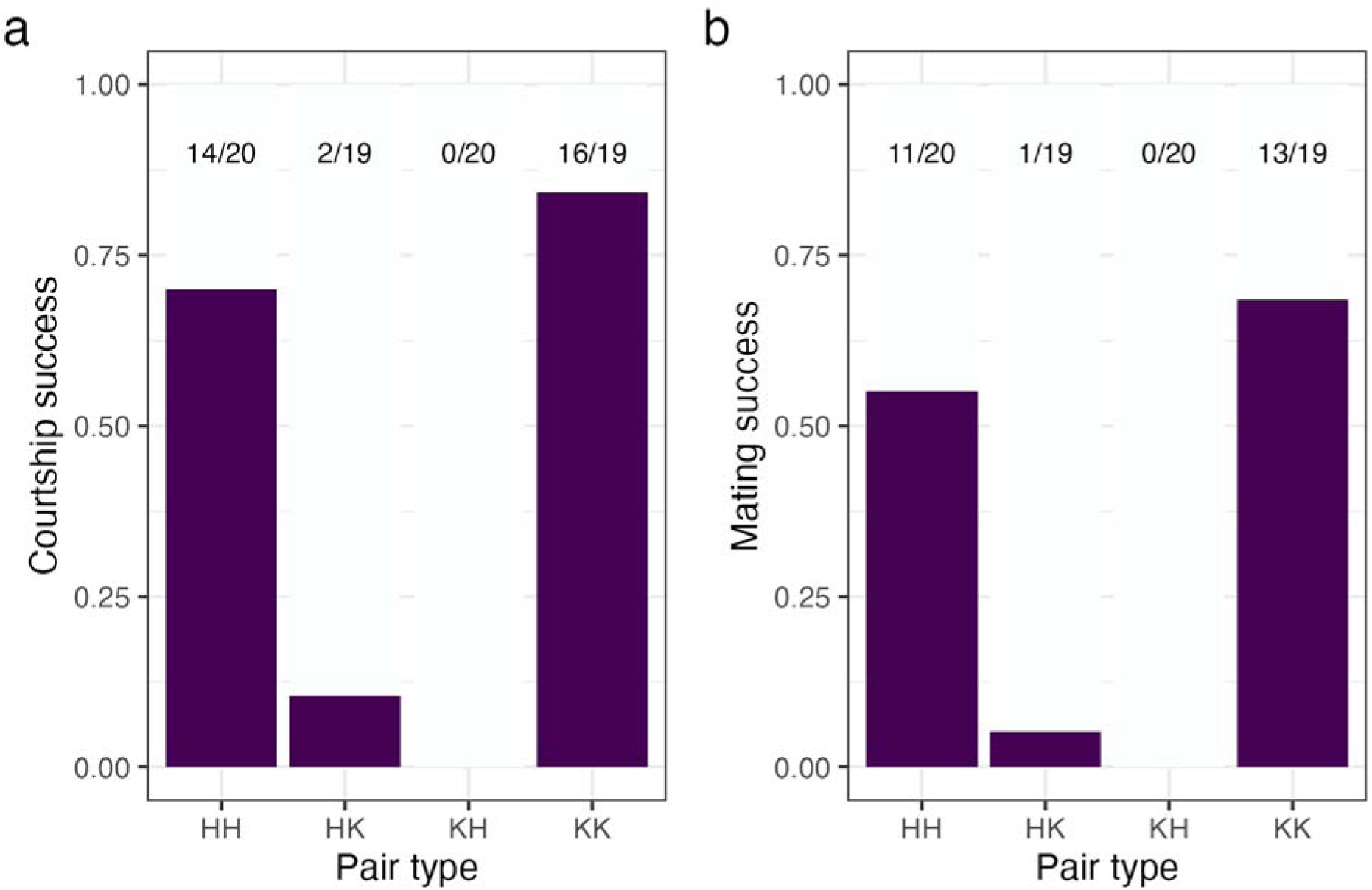
Courtship (a) and mating (b) success (percentage of pairs where a micro and macro were transferred, respectively) in interspecies mating trials of *L. hualalai* and *L. kona*. The four possible mating combinations are: HH: conspecific *L. hualalai* pair, HK: male *L. hualalai* paired with a female *L. kona*, KH: male *L. kona* paired with a female *L. hualalai* and KK: conspecific *L. kona* pair. Note that these are not independent trials: the same individual was first tested in a heterospecific contest and then paired with a conspecific.

## 4. Discussion

Understanding the role of different reproductive isolating barriers which keep closely related species from hybridizing, is an important goal in evolutionary biology. Habitat and sexual isolation are often the first barriers between closely related species, but they are seldom studied together, begging the question if either barrier is sufficient or more important in maintaining reproductive isolation on secondary contact. Through a combination of field observations and lab experiments, we did the first comprehensive study of habitat and sexual isolation in the rapidly speciating crickets of the genus *Laupala*. Using genomic data we first confirmed that sympatric species *L. hualalai* and *L. kona* are genetically distinct, a pattern you can only observe under strong reproductive isolation. Next, we probed what factors are maintaining these species barriers. We measured different aspects of their microhabitats in the wild, used isotope proxies to infer diet differences, monitored singing activity, documented sexual trait variation in the lab, measured female preferences and conducted inter-species mating trials in the lab. Our results indicate that despite broad ecological equivalence of their habitats (with the possible exception of diets), sexual isolation is sufficient to keep species boundaries intact between *L. hualalai* and *L. kona* in their natural habitats of Hawaiian rainforests.

Before we delved into what factors kept species boundaries intact, we wanted to confirm that *L. hualalai* and *L. kona* were reproductively isolated for a prolonged period of time. We inferred this from genetic data: strong reproductive isolation would keep gene pools isolated and distinct, weak reproductive isolation would cause interspecies mating which would be reflected as extensive admixture in the genetic data. Our genomic PCA and ADMIXTURE analyses confirm that *L. hualalai* and *L. kona* are genetically distinct populations with only a few *L. kona* individuals showing very low levels of admixture. The maximum likelihood phylogeny also classifies the two species as separate monophyletic groups. Previous phylogenetic analyses (Mendelson & Shaw, 2005) showed that they evolved in allopatry and then moved into their current sympatric range. Our genomic data shows that *L. hualalai* and *L. kona* have maintained species boundaries while in sympatry. We next asked whether these strong reproductive barriers are due to preferences for diverse habitats or divergent mating phenotypes.

Habitat isolation can reduce the encounter frequency of two species when they occupy different parts of their ecological niche. However, our data show broad ecological equivalence: crickets of both *L. hualalai* and *L. kona* occupy the same vegetation, from similar elevation from the ground and micro-habitats with similar substrate temperatures. In fact, most crickets of either species were mostly found on the ground, hiding in the leaf litter. This is in a slight contrast to an earlier study (Xu & Shaw, 2020) on sympatric *L.cerasina* and *L.pruna* where the crickets called from significantly different heights, although with a huge overlap in spatial distribution. This disparity may be explained by the time between the colonization of two species. In contrast to the recently diverged *L. hualalai* and *L. kona* (∼0.5 Mya), *L.cerasina* and *L.pruna* belong to more distantly related clades (∼3.7 Mya), indicating that the latter pair probably had more time to evolve some form of ecological differences, whereas the former pair did not. One limitation of our study is that we can only study singing males, it could be the case that the two species share singing sites but otherwise partition their micro-habitats, a hypothesis that can be tested in future studies. The nearest neighbor distances (NND) between conspecific and heterospecifics implies that *L. hualalai* and *L. kona* form mixed-species aggregations, in line with previous work in *L.cerasina* and *L.pruna* (Xu & Shaw, 2020). This spatial mixing creates ample opportunity for reproductive encounters between species, meaning something other than habitat isolation is maintaining species boundaries.

However, the species did differ in their δ^13^C values. The δ^13^C values from animal tissues indicate dietary carbon sources allowing distinguishing between plant and animal sources, C3 or C4 plants, among other distinctions (DeNiro & Epstein, 1978). δ^15^N values, on the other hand, reflect the animal’s trophic position in the food web (Boecklen et al., 2011; Minagawa & Wada, 1984). From our results of distinct δ^13^C profiles between *L. hualalai* and *L. kona* but similar δ^15^N profiles, we can speculate that they probably have different dietary sources while having a similar trophic position on the Hawaiian rainforest food webs. However, in the absence of direct behavioral observations of feeding behavior across both species or more precise methods like gut content analysis, it is difficult to say for sure if *L. hualalai* and *L. kona* have different diets. Different dietary sources can in theory reduce interspecies encounters if they have different foraging sites, under the assumption that they can forage and mate at similar times. But our NND data shows that singing males form mixed species aggregations and singing is an important starting point in *Lauapal* mating rituals, making it safe to assume that there is enough opportunity for interspecies encounters during their mating periods. However, nutrients in the diet can indirectly affect sexual behavior by affecting cuticular hydrocarbon profile expression in insects (Claudio-Piedras et al., 2021; Ng et al., 2018; van Wilgenburg et al., 2022), which is an important sexual phenotype and divergent between *L. hualalai* and *L. kona* (see Results 3.4).

Divergence in timing of reproductive events may contribute to the maintenance of species boundaries and coexistence of closely related species (Coyne & Orr, 2004). For example, divergence in the emergence time and reproductive period of periodic cicadas partly contributes to their speciation (Cooley et al., 2003; Grant, 2005; Ritchie, 2001). Divergence of reproductive behavior can happen at annual, seasonal or diurnal scales. In *Laupala* and in other crickets, males sing to attract females, so we quantified male singing activity as a proxy for reproductive timing. We found that peak singing activity of *L. hualalai* (9-10 am) is earlier than *L. kona* (1-3 pm), which can reduce interspecies mating encounters. Daily variation in reproductive behavior may contribute to reproductive isolation as has been shown in aphids (Abbot & Withgott, 2004), moths (Silvegren et al., 2005) and water-fleas (Wen Deng, 1997). This asynchrony in singing activity in *Laupala* could be a result of reproductive character displacement caused due to interactions between *L. hualalai* and *L. kona*. Whether this asynchrony in male singing behavior can act as an effective mating barrier depends on if females have a corresponding pattern of female receptivity. In either case, asynchrony in diel singing activity will not be a complete reproductive barrier between *L. hualalai* and *L. kona*, as there is some overlap of their activity patterns (see **Figure 5**). We investigated other components of sexual isolation, which could further reduce interspecies matings.

The primary axis of phenotypic divergence in the *Laupala* genus is the pulse rate of the male song. We show that *L. hualalai* and *L. kona* males sing with significantly different pulse rates and this divergence is documented both in the wild and in laboratory settings. Songs collected in the lab had higher pulse rates than songs collected in the wild, because their natural habitat was often colder (due to high elevation of the cloud forest and frequent rainfall) than a controlled laboratory setting, and we know pulse rates increase with air temperature (unpublished data). *Laupala* species have narrow variation in species-specific pulse rates. Amongst our extensive sampling, both *L. hualalai* and *L. kona* populations have typical levels of variation as that of other species, and we did not sample any individuals with intermediate pulse rate values. In lab crosses, F1 and backcrossed hybrids possess intermediate pulse rates between their parental species (Shaw, 1996). The lack of phenotypic intermediates at our study sites suggests that species boundaries are strong and no contemporary hybrids exist. Females always preferred conspecific songs over heterospecific songs in a phonotaxis arena, indicating strong preference for conspecific pulse rate. This confirms years of work on conspecific pulse rate preferences in *Laupala* females (Grace & Shaw, 2011, 2012; Mendelson & Shaw, 2002; Shaw, 2000) and supports the hypothesis of genomic coupling of traits and preferences as documented in a different Laupala species pair (Shaw & Lesnick, 2009; Wiley & Shaw, 2010; Xu & Shaw, 2021). This suggests that in the wild, when a female *Laupala* hears two different songs, it predominantly approaches the conspecific male.

A typical courtship ritual in *Laupala* begins with a female approaching a singing male, followed by extended periods of antennal interactions (Shaw & Khine, 2004). During these interactions, crickets assess the cuticular hydrocarbons on their potential mate. We found substantial species variation in CHC profiles of *L. hualalai* and *L. kona*. CHCs are a complex multivariate phenotype, so we summarized the variation in a principal components analysis, which clustered the two species separately. In *Laupala*, males elicit separate responses to female and male CHCs (Stamps & Shaw, 2019) and species in the genus have distinct CHC profiles (Mullen et al., 2007), suggesting that crickets use this information for species-specific mate discrimination, although this hypothesis needs additional testing. However, there is ample evidence of mate discrimination using CHCs in field crickets (Tyler et al., 2015), sagebrush crickets (Steiger et al., 2013), *Drosophila* flies (Shahandeh et al., 2018), beetles (Xue et al., 2016; Zhang et al., 2014) and butterflies (Ômura et al., 2020).

Finally, we did a direct test of sexual isolation: an inter-species mating (in)compatibility experiment which clearly revealed that there is complete sexual isolation between *L. hualalai* and *L. kona*. We performed no-choice mating trials, which normally suffers from the limitation of under-estimating mating compatibility, because the fitness consequences of not mating can be huge so individuals may choose an unsuitable mate. However, this limitation makes our case stronger: in-spite of being in a no-choice scenario, both *L. hualalai* and *L. kona* only courted and mated with conspecifics. *Laupala* crickets have a long and protracted courtship ritual lasting 5-7 hrs involving the male producing and transferring many spermless micros followed by the sperm filled macros (Shaw & Khine, 2004). A male does not produce a macro if it has not successfully transferred a few micros. We find that the courtship ritual between heterospecific pairs gets stalled early on, with very few males successfully transferring a micro to a heterospecific female (only 2/37 did). Only one male (out of 37) ever produced a sperm filled macro and transferred to the heterospecific female, showing very strong sexual isolation. On the other hand, these males readily mated conspecific females when presented with them 1-2 days after interacting with the heterospecific female.

Our results together show that despite the ample possibility of reproductive encounters between *L. hualalai* and *L. kona* due to them occupying the same micro-habitats and existing in mixed-species aggregations, they do not mate with each other. We comprehensively catalogued different components of habitat and sexual isolation and showed that sexual isolation is sufficient to keep closely related *Laupala* species reproductively isolated in sympatry. Since sexual barriers are one of the early acting barriers (after habitat barriers) when different species interact, it also appears to be the strongest barrier to gene flow in our system, reducing the need for any later acting pre- or post-zygotic barriers. This result of sexual isolation being the strongest barrier in sympatry is in line with many studies in seed plants (Christie et al., 2022), fungi (Gac & Giraud, 2008), *Drosophila* flies (Yukilevich, 2012) and *Heliconius* butterflies (Garzón-Orduña & Brower, 2018; Mérot et al., 2017; Rosser et al., 2019), with the caveat that not all studies have characterized the ecological barriers. Our study provides rare empirical evidence that shows that sexual isolation can evolve and persist without habitat isolation. This is in contrast to the general conclusions from most theoretical studies (Chesson, 2000; Germain et al., 2021; Servedio & Bürger, 2014) which conclude that some degree of ecological differentiation is needed for coexistence of species. We advocate for more studies of both habitat and sexual isolation in other systems to understand the complex interaction between habitat and sexual isolation in preventing gene flow in sympatry.

## Acknowledgements

We are grateful to Natalie Graham and Patrick Hart for providing lab resources and office space while conducting fieldwork on the Big Island of Hawaii. We thank the “Lunch Bunch” discussion group in the Department of Neurobiology and Behavior at Cornell University for fruitful discussions on the project. We thank Quinn Tonole for help with the behavioral trials and Ella Zhao for help with animal care. We also thank the University of Wisconsin Biotechnology Center DNA Sequencing Facility (Research Resource Identifier – RRID:SCR_017759) for providing GBS sequencing services.

## Data and Code availability

All data and R codes will be made publicly available upon publication of the manuscript.

